# Enhancing generating and collecting efficiency of single particle upconverting luminescence at low-level power excitation

**DOI:** 10.1101/2019.12.17.879858

**Authors:** Chenshuo Ma, Chunyan Shan, Kevin Park, Aaron T. Mok, Xusan Yang

## Abstract

Upconverting luminescent nanoparticles are photostable, non-blinking, and low chemically toxic fluorophores that are emerging as promising fluorescent probe at single-molecule level. High luminescence intensity upconversion nanoparticles (UCNPs) is achieved with highly doped rare-earth ions co-doped (20% Yb^3+^) using high excitation power (>2.5 MW/cm^2^). However, such particles are inadequate for *in-vitro* live-cell imaging and single-particle tracking since high excitation power can cause photodamage. Here, we compared UCNPs luminescence intensities with different dopants concentrations and presented a more efficient (∼7x) UCNPs at low excitation power by increasing the concentrations of Yb^3+^ and Tm^3+^ dopants (NaYF_4_: 60% Yb^3+^, 8% Tm^3+^) and adding a core-shell structure.

## Introduction

Upconversion nanoparticles (UCNPs) possess the ability to convert low-energy infrared photons into high-energy visible or ultraviolet photons^1^. This unique anti-Stokes feature has enabled various applications over the past decade, spanning from solar energy harvesting, super-resolution microscopy, anti-counterfeit luminescent imprinting to agricultural crop care^2–5^. Among the various nanoparticle systems developed thus far, semiconductor nanocrystals or quantum dots (QDs) are widely used^6,7^. UCNPs typically consist of a NaYF_4_ crystalline host matrix doped with rare-earth-based lanthanide sensitizer and activator. Such particles can be designed and synthesized base on desirable physicochemical characteristics such as sizes, shapes, optical properties and magnetic properties^8–11^.

UCNPs exhibit neither photo blinking on millisecond and second-time scales nor photobleaching even with hours of continuous excitation. In addition, the nanoparticles feature efficient cellular uptake while retaining low cytotoxicity and display high optical penetrating power in deep tissue with minimal background noise^12,13^. As a result, UCNPs is becoming one of the most promising nanoparticle systems for *in-vivo* biological imaging, bio-sensing, and nanomedicine, with continuing efforts to improve their properties by designing new synthetic strategies^14^.

Single nanoparticle imaging is essential for cell tracking. Studies include long-term tracking of UCNPs in living cells^15^, deep sub-diffraction fluorescence microscopy^16^, single-molecule tracking under lower irradiance^17^, using agarose-gel electrophoresis to make background-free labels^18^ and improving upconverting fluorescence with natural bio-microlens^19^. Single nanoparticle imaging is essential for the detection of single-cell imaging in techniques involving cellular tracking and providing accurate cellular movement.

For bioimaging applications *in*-*vitro*, high excitation power is not recommended. Imaging and logitudinal real-time tracking of living cells prefer low power excitation. For example, power dependent analysis are often used in various studies including investigating energy transference relations between dopants^20^, economically feasible UCNPs in anti-counterfeiting ink printing^21^, thin-film synthesis utilizing electro-deposition^22^, and upconverting luminescence depletion phenomena via simultaneous wavelength excitation^23^. Therefore, in the practical usage of UCNPs in bio-informatics, maximizing luminescence efficiencies is of great significance. In this article, emphasis is placed on such augmentation of upconversion intensity in response to power irradiance.

Plenty of studies had aim to improve upconverting luminescence in the advancement of synthesis methodology and instrumental characterization at nanoscale over decades. Enhancement strategies include optimizing crystal hosts and dopant concentrations, coating inert/active shells around UCNP, tuning UCNPs’ sizes, and coupling UCNPs with plasmonic noble metal nanostructures. In the synthesis of UCNPs, a higher dopant concentration of sensitizer ions will enhance luminescence. However, an oversaturation of sensitizer concentration concerning activator concentration will decrease luminescence^24^. This concentration quenching consequence is an important factor to take into consideration when addressing the dopant concentrations in such particles^25,26^. The addition of an inert/active shell as a core-shell arrangement of particles provides a protective layer that prevents luminescence quenching associated with surface defects on the core particles, but at a cost of increasing nanoparticle size^27^. It is found that typical lanthanide-based UCNPs containing smaller particles below ten nanometers in size endure more difficulty in displaying efficient upconverting luminescence due to their cubic crystal lattice structure^28^. However, nanoparticle size for efficient cellular uptake favors smaller sizes, which can decrease the toxicity of the particles within the subcellular regions and increase bio-compatibility^29,30^.

There are several reports on acquiring high contrast cellular imaging thorough testing with a customized confocal microscope with a moderate resolution, in response to the saturation effect of upconverting luminescence^31^. The saturated effect is easily achieved with high N.A. confocal microscopy with a power density of about 10 KW^−1^ MW/cm^2^ while the nanoparticle have a saturated threshold of 2.5 MW/cm^2 32^. For the living cell imaging, a probe with high quantum efficiency and low saturated excitation threshold power and will benefit long term living cell imaging or single-particle tracking.

In this article, multifarious methods to improve luminescence intensity in UCNPs are investigated. A more power efficient UCNPs are synthesized by scrutinizing dopant concentration, particle size and core-shell influence. The luminescence efficiency variation by changing each of these attributes are also quantified and compared. The benefits each particular attribute provides with respect to emission efficiency are quantified and compared to one another. The numerical factors at which the luminescence is increased in better performing nanoparticles in regard to the maximum power densities recorded for lesser optimal nanoparticles are emphasized. We explored UCNPs with highly doped Yb^3+^ sensitizer ions and Tm^3+^ activator dopant in a NaYF_4_ host and their core-shell structures. This combination of dopants are typical lanthanide dopants used in UCNPs research and have been used in bio-informatics investigations^33,34^. With a dopant concentration of 60% Yb^3+^, 8% Tm^3+^ in NaYF_4_ host, we present a ∼7x more efficient UCNPs that will be well-suited for applications in live cell imaging.

## Results and discussion

There are various methods to increase the upconverting luminescence with concentrated samples. We also found multiple methods to increase the upconverting luminescence for single nanoparticle^26,35^, as shown in Figure 1. The nanoparticles’ size is an important aspect which can influence the upconverting luminescence, as different nanoparticle sizes adjust the surface/volume ratio and directly influences the defects/unit. The size of the nanoparticles is controlled below around 50 nm in consideration towards use for bio-applications. We use optimized synthesis methods to synthesize the individual nanoparticles within similar sizes, as shown in Figure S1 and S2. Similar sizes will maintain the same surface/volume ratio, which will greatly eliminate of the influence of surface defects.

**Figure 1.**
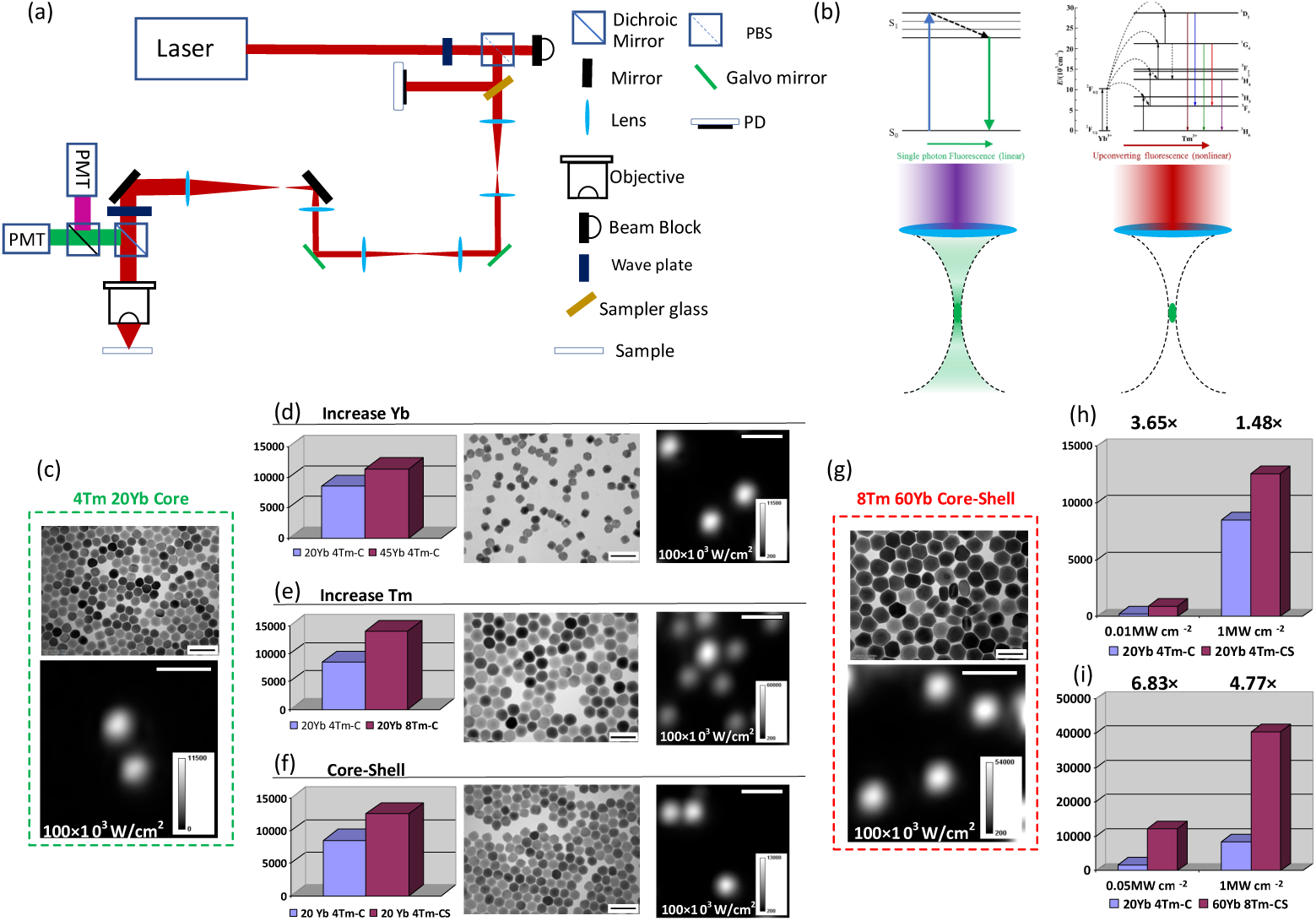
(a): Schematic illustration of the system setup for customized laser scanning microscopy; The excitation source is a 980nm continue wave (CW) laser (power density of 30 MW/cm^2^) which is focused on the sample through an oil immersion objective 100× objective lens (NA 1.4, Olympus). The emission from the sample is reflected by the dichroic mirror into the detector. A large field PMT (Multialkali Amplified PMT, PMT1001, Thorlabs) detector is employed to collect the emission fluorescence photons. The scanning is achieved by two conjugated placed galvanometer mirrors. A motorized wave plate and polarizing beam splitter (PBS) are employed to control the power, and a cover glass and photon diode (PD) are utilized to sample the power on the sample after objective. (b): Spatial confinement of generated fluorescence generation with upconverting nonlinear excitation. In one photon fluorescence process the whole illumination cone will generate fluorescence signal, but upconverting nonlinear generated signal is only localized the focal spot. Upconverting nonlinear process generated a fluorescence photon with absorbing > 2 photons, whereas one photon fluorescence process case absorbs 1 photon. Transmission electron microscopy (TEM) images and microscopy quantitative measurement of whole upconverting spectrum luminescence emission of single UCNPs of (c): NaYF_4_: 4% Tm^3+^, 20% Yb^3+^ UCNPs; (d): NaYF_4_: 4% Tm^3+^, 45% Yb^3+^ UCNPs; (e): NaYF_4_: 8% Tm^3+^, 20% Yb^3+^ UCNPs; (f): NaYF_4_: 4%Tm^3+^, 20% Yb^3+^@NaYF_4_ UCNPs; (g): NaYF_4_: 8% Tm^3+^, 60% Yb^3+^@NaYF_4_ UCNPs, scale bar is 100nm. Comparison of integrated single-particle upconverting luminescence emissions of (h): core (4% Tm^3+^, 20% Yb^3+^) and core–shell; (i): NaYF_4_: 4% Tm^3+^, 20% Yb^3+^ vs. NaYF_4_: 8% Tm^3+^, 60% Yb^3+^@NaYF_4_ UCNPs under different excitation power. Scale bar is 1um, pixel dwell time, 200 µs.

Since Yb^3+^ acts as sensitizer that can transfer the 980 nm phonon energy to rare-earth ions. It is assumed that the high sensitizer concentration combined with high rare-earth ions doped UCNPs will be much brighter under high excitation power. Shown in Figure 1(d), 45% Yb^3+^ doped UCNP is 1.33 times higher than the 20% Yb^3+^ (Figure 1(c)) doped particles under 1 MW cm^−2^ power density. Increasing Yb^3+^ concentration will decrease the atomic distance from sensitizers to activators, thus, energy transfer will be more efficient by such decrease in ions distance when high excitation power continually transfers energy from Yb^3+^ to activators. The luminesce intensity of single nanoparticle with different dopant concentration has been measured by using a customized laser-scanning microscope (shown in Figure 1(a)). The single nanoparticle’s luminesce intensity is represented as the maximum pixel value for each Gaussian spot.

Based on the previous report, a brighter upconverting luminescence can be obtained under high excitation power in high activator concentration (4% Tm^3+^) co-doped with 20% Yb^3+^ UCNPs^32^. We tried to use a higher activator concentration to further increase the luminescence, which is 8% Tm^3+^ doped UCNPs. As shown in Figure 1(e), the same trend occurs such that the luminescence enhances 1.64 times higher.

Concentration quenching also can be overcome at high excitation powers by introducing a thin layer of an inert shell. From Figure 1(f), the core-shell particle increases the luminescence by 1.48 times compared with the exposed particles. Yb^3+^-Yb^3+^ energy migration to surface defects, which lead to quenching, is primarily responsible for depopulation of the core UCNPs. This result demonstrates that at single-particle laser powers, the quenching typically associated with high Yb^3+^ content can be suppressed by the external inert shells.

At the same time, the Tm^3+^ mainly produce ^3^H_4_→^3^H_6_ within low excitation power, but a different situation appears when excited under high power. As shown in Figure 1(h), the blue wavelength (^1^D_2_→^3^F_4_, ^1^G_4_→^3^H_6_), red wavelength (^1^G_4_→^3^F_4_) and near-infrared (^1^D_2_→^3^F_2_, 1D_2_→^3^F_3_, ^3^H_4_→^3^H_6_) are all verified in Figure 1(b). From the different wavelength emission changes under different excitation power, we can find out that the electron population on ^1^D_2_ and ^1^G_4_ states increased faster than ^3^H_4_ state when increasing excitation power. The ^3^H_4_ state release the 800 nm emission, however, when additional emission energy is transferred from Yb^3+^ under high excitation power, the ^3^H_4_ state eventually achieves full capacity and is excited to higher ^1^D_2_ and ^1^G_4_ states.

It is worth noting that the increasing luminescence intensity ratio between core-shell and core only UCNPs is higher under 10 kW cm^−2^ (3.65 times) than 1 MW cm^−2^ (1.48 times), as shown in Figure 1(h). Low power density is required for *in-vivo* imaging; therefore, importance is placed on attaining the same luminescence intensity with a lower excitation power density. Based on all the listed luminescence enhancement methods, such intensity with low power density is produced by using higher sensitizer and activator concentration combinations along with a core-shell structure. As Figure 1(i) shows, the core-shell structure with high dopant concentrations (8% Tm^3+^, 60% Yb^3+^, shown in Figure 1(g).) is 6.83 times higher than the normal UCNP particles under 0.05 MW cm^−2^.

To systematically compare the brightness of all the improved methods, we measured the power-dependent luminescence curves for single particles from 5 kW cm^−2^ to 10 MW cm^−2^, as shown in Figure 2 (a)-(h). The quantitative intensity images are shown in Figure 2 (i)-(l) and every image is taken within the same area for each sample. From the previous recordings of the data obtained from the sensitizer concentration and power dependence testing, it is foreseen that the luminescence would be enhanced with the increase in excitation power. From Figure 2(a) and (d), which are the 4% Tm^3+^ doped UCNPs, it shows the higher sensitizer concentration will not only increase the luminescence intensity under each excitation power, but also can shift the power dependence curve to lower power. In other words, we can use lower power excitation with higher sensitizer concentration UCNPs to get the same intensity as those using high power excitation with lower sensitizer concentration. Figure 2(b) and (f), 8% Tm^3+^ doped UCNPs, shows the same trends as 4% Tm^3+^ particles. Although, higher Tm^3+^ dopant concentrations will increase the luminescence intensity under each excitation power.

**Figure 2.**
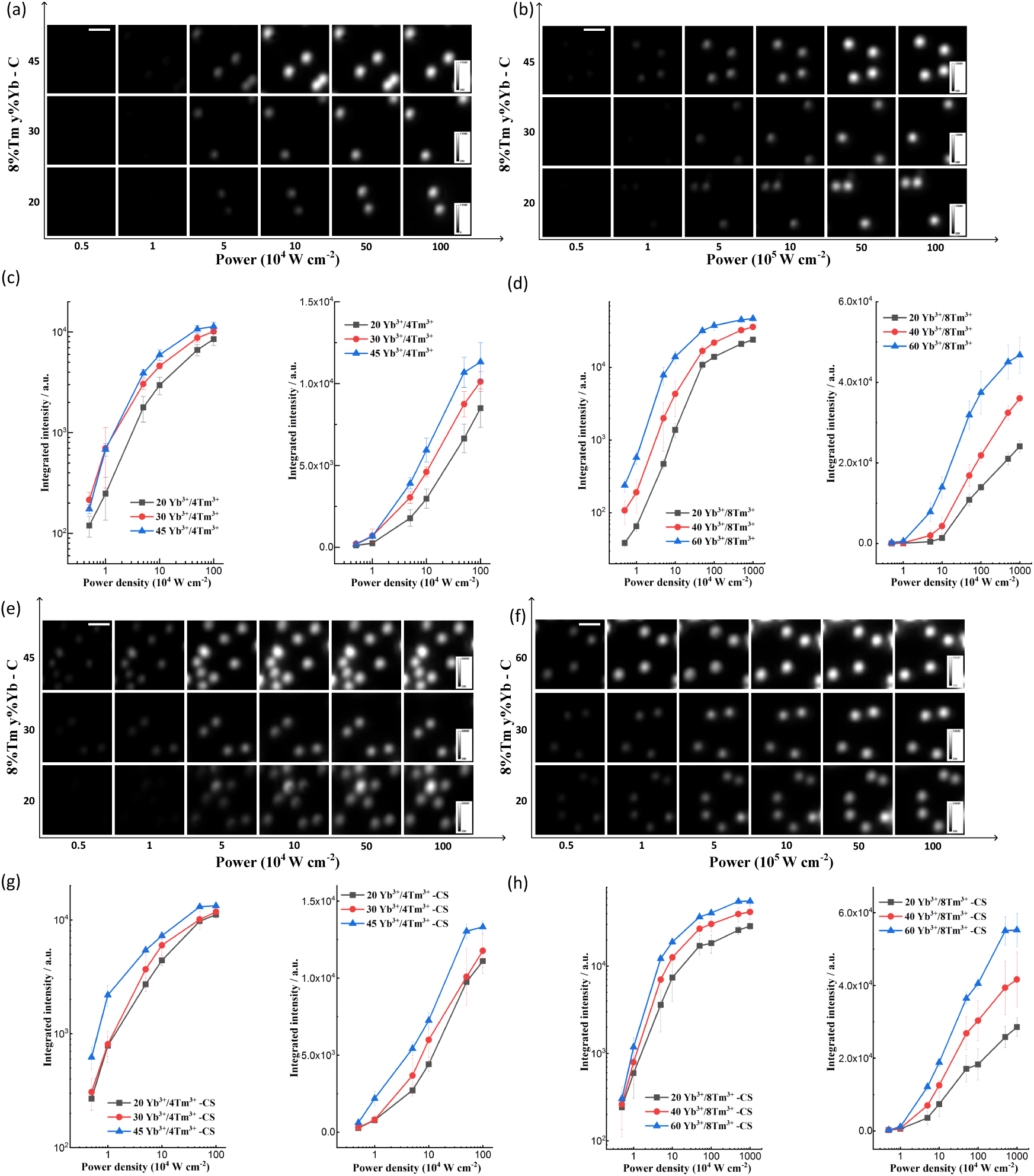
Microscopy quantitative measurement of whole upconverting spectrum luminescence emission of (a)single 4% Tm^3+^ UCNPs vs. Yb^3+^ concentration, (b)single 8% Tm^3+^ UCNPs vs. Yb^3+^ concentration. (e)single 4% Tm^3+^ core-shell UCNPs vs. Yb^3+^ concentration, (f)single 8% Tm^3+^ core-shell UCNPs vs. Yb^3+^ concentration. The power dependent saturation curves of (c)single 4% Tm^3+^ UCNPs vs. Yb^3+^ concentration, (d)single 8% Tm^3+^ UCNPs vs. Yb^3+^ concentration, (g)single 4% Tm^3+^ core-shell UCNPs vs. Yb^3+^ concentration, (h)single 8% Tm^3+^ core-shell UCNPs vs. Yb^3+^ concentration. The saturation curves under different power densities from 5 kW cm^−2^ to 10 MW cm^−2^ obtained with large field PMT detector-based laser scanning microscopy. A log scale is used for the y axis in the left panels of (c)-(d) and (g)-(h), and a linear y scale is employed in the right panels of (c)-(d) and (g)-(h). Scale bar is 1um; pixel dwell time, 200 µs. Error bars represent standard deviation from the mean value.

Another remarkable phenomenon is the power dependence slope increase by introducing a core-shell structure, in which surface defects are decreased. We can use 0.1 MW cm^−2^ for 4% Tm^3+^/45% Yb^3+^ core-shell particles to obtain the same highest luminescence intensity of the related core particle under 0.01 MW cm^−2^. In comparing the 4% Tm^3+^ UCNPs, with 8% Tm^3+^ doped particles, the power dependence slope for 8% Tm^3+^ is higher than 4% Tm^3+^ UCNPs. This indicates that the 8% Tm^3+^ doped UCNP can reach its saturation point at a much lower power than 4% Tm^3+^ UCNP. The 8% Tm^3+^ doped UCNP is able to reach to the same luminescence intensity by using much lower excitation power. For instance, the highest luminescence intensity of NaYF_4_: 4% Tm^3+^/20% Yb^3+^@ NaYF_4_ UCNP is under 1 MW cm^−2^, the same intensity can be reached under 0.02 MW cm^−2^ by the NaYF_4_: 8% Tm^3+^/60% Yb^3+^ @ NaYF_4_ UCNP. Based on these results, the excitation power can decrease by more than one order of magnitude by using high Tm^3+^ with high Yb^3+^ dopants. It is indicative that the short distances between sensitizer and activator ions as well as the high-power excitations that increase the electron populations of excited states, play an important role in protecting the surface quenching of same sized UCNPs.

The spatial resolution and collection efficiency advantage of the large field PMT detector-based laser scanning microscopy for high nonlinear upconverting fluorescence nanoparticles. Several groups have reported their results of single upconverting nanoparticle characterization employing confocal microscopy^17,36–39^ as well as confocal-based laser scanning sub-diffraction microscopy^3,16,40^. The merits of the nonlinear contrast mechanisms are that the signal (S) depends supralinear (S∝I^n^) to the excitation light intensity (I). As a result, when focusing the laser beam through a microscope objective, high order nonlinearity absorption is spatially confined to the perifocal region. The high order non-linear property of upconverting luminescence demoed in Figure 2 can benefit not only the spatial resolution of laser scanning microscopy but also the out-of-focus background could be a suppressed by high order nonlinear generated fluorescence. Thus, pinhole is not needed to reject out-of-focus background. As important as the maximization of signal generation is the optimization of collection and detection efficiencies. The pinhole will also degrade the collection efficiency of the generated fluorescence once it is not well coupled. High-order nonlinear microscopes (Figure 1 (a)) use photomultiplier tubes, available with large sensitive areas and reasonable quantum efficiencies rather than the pinhole based-detector, also coming with a high spatial resolution of nonlinear upconverting fluorescence nanoparticle.

The resolution of confocal images degrades as the laser power increases because the excitation power has exceeded the nonlinear threshold. For microscopy, the image of any object is acquired by convolving the point spread function (PSF) of the optical system^41^. For confocal microscopy, *PSF*_*confocal*_ = *PSF*_*illumination*_ × *PSF*_*detection*_. For N-th order nonlinearity laser scanning microscopy with whole area detection the effective PSF, *PSF*_*N*−*order*_ = (*PSF*_*illumination*_)^*N*^ = *PSF*^*N*^ (*u, v*)^41^. As demoed in Figure 2, the log scale power dependence fluorescence saturated curves have a higher slope under a low power density. For example, 4% Tm^3+^, 20% Yb^3+^ core UCNPs have a slope of ∼1.6 below 1 MW cm^−2^ (saturated Intensity), and 8% Tm^3+^, 60% Yb^3+^ core-shell UCNPs have a slope of ∼2 with below the saturated intensity. In Figure 3, the image under lower power density of 5 kW cm^−2^ have a much smaller full width at half maximum (FWHM), 279.0 nm, comparing to the case with 100 kW cm^−2^ (saturated absorbed). Note that 4% Tm^3+^, 20% Yb^3+^ core UCNPs have a much narrowed PSF, but really low efficiency of generating fluorescence under low density excitation power. However, the 8% Tm^3+^, 60% Yb^3+^ core-shell UCNPs have also an enhanced resolution, 294.4 nm, due to the high order nonlinearity of upconverting fluorescence with decent fluorescence generating efficiency under low power excitation density (5 kW cm^−2^).

**Figure 3.**
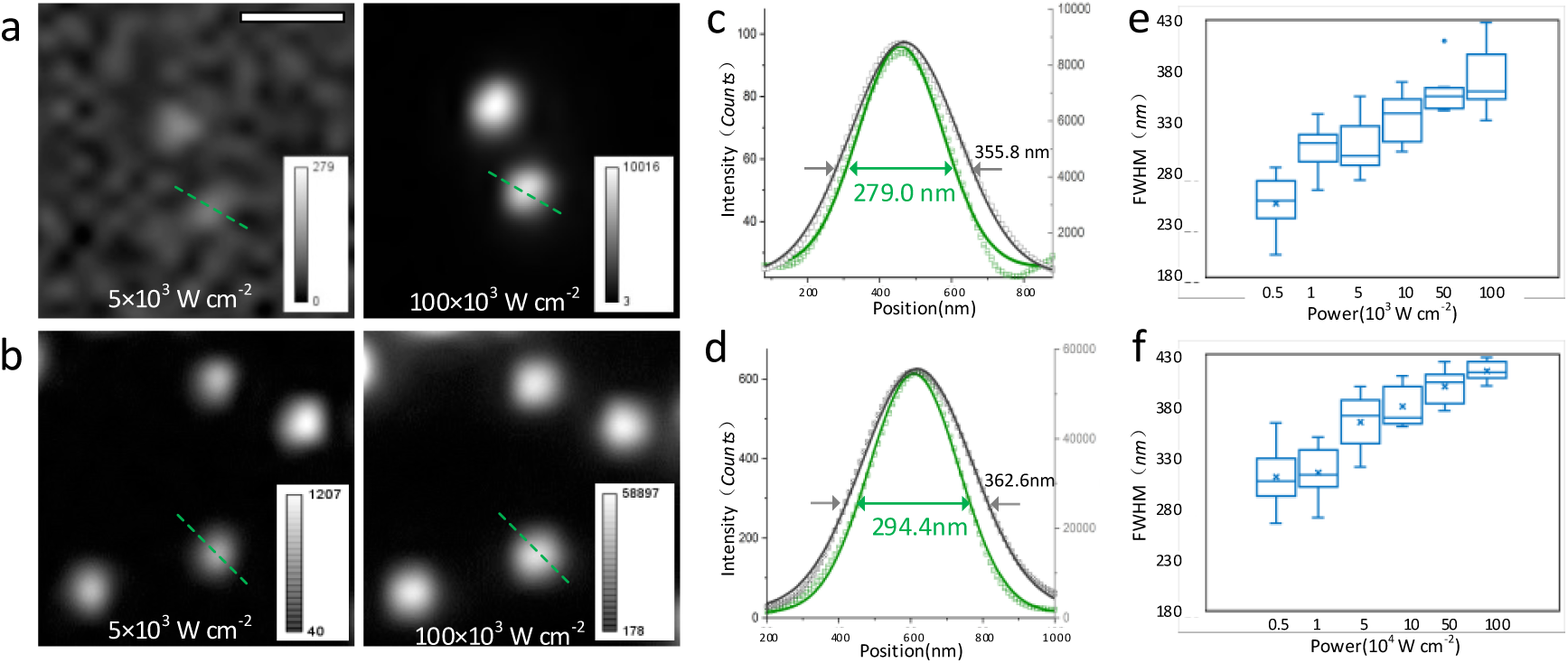
Laser scanning upconverting nonlinear image and saturated image of single 4% Tm^3+^, 20% Yb^3+^ core UCNPs and single 8% Tm^3+^, 60% Yb^3+^ core-shell UCNPs. (a) 4% Tm^3+^, 20% Yb^3+^ core single UCNPs under nonlinear excitation (5×10^3^ W cm^−2^) and saturated excitation (100×10^3^ W cm^−2^), respectively. (b) 8% Tm^3+^, 60% Yb^3+^ core-shell single UCNPs under nonlinear excitation (5×10^3^ W cm^−2^) and saturated excitation (100×10^3^ W cm^− 2^), respectively. The cross-section on the green dashed line in (a) and (b) are shown in (c) and (d), the fitted curves of nonlinear excitation (green) have a smaller FWHM then saturated excitation ones (in gray) due to different order of non-linearity. (e) and (f) are the FWHM under different power density excitation of 4% Tm^3+^, 20% Yb^3+^ core UCNPs and 8% Tm^3+^, 60% Yb^3+^ core-shell UCNPs, respectively.

Using UCNPs with continuing wave (CW) infrared excitation has similar merits as a multiphoton microscopy for deep tissue brain imaging, longer scattering length, and high order nonlinearity. As the microscopy focus goes deep, the benefits of confocal detection diminish, and attention has to be paid to the detection of scattered light. The scattered light will spread to a volume that is much larger than when the excitation focus is deep below the scattering tissue. The large field detector-based laser scanning microscopy using nonlinear upconverting nanoprobe have the ability to realize deep tissue imaging which is similar to fs ultrafast laser-based multiphoton microscopy. By using upconverting microscopy with a CW laser, it will bring nonlinear deep tissue microscopy more cost efficient and portable.

## Conclusion

This work emphasizes the importance of controlling the activator and sensitizer dopant concentration to decrease the excitation power for emitting the same upconverting luminescence intensity. We found the optimal activator and sensitizer concentration in single UCNPs that can increase the upconverting luminescence and increase the power dependence slopes. Inserting shell coated to passivate the quenchers on the surface will shift the power dependence curve to lower excitation power. Also, higher sensitizer concentration will help to shifts the power dependence curve to a lower excitation power. The core-shell structure will help to improve the luminescence intensity. The particle’s luminescence intensity will be easier to reach to the saturation intensity under lower power when higher activator concentration.

## Acknowledgments

C.S thanks the support by Maintenance Fund of National center of Protein Science, Peking University.

