## Supplementary material for "Enhancing generating and collecting efficiency of single particle upconverting luminescence at low-level power excitation": Supplemantary methods and EM figures

### Experiment:

#### 1. Materials

$YCl_3·6H_2O$ (99.99%), $YbCl_3·6H_2O$ (99.99%), $TmCl_3·6H_2O$ (99.99%), $NH_4F$ (>98%), NaOH (>97%), Oleic Acid (OA, 90%), and 1-Octadecene (ODE, 99%) were purchased from Sigma-Aldrich and used as received without further purification.

#### 2. Synthesis of hexagonal-phase $NaYF_4$ nanoparticles:

$NaYF_4$ nanoparticles were synthesized using a typical method as previous work. Typically, a methanol solution of 0.04 mmol $TmCl_3·6H_2O$, 0.2 mmol $YbCl_3·6H_2O$, 0.76 mmol $YCl_3·6H_2O$ were mixed with 6 ml OA and 15 mL ODE in a 50 mL round bottom flask. The mixed solution was heated up to 150˚C for 30 min until it became clear. With the gentle flow of argon gas through the reaction flask, the solution cooled slowly to room temperature. Methanol solution dissolved with 4mmol $NH_4F$ and 2.5 mmol NaOH was added to the flask with vigorous stirring for more than 30 min. Then, the mixed solution was heated up to 90˚C to evaporate methanol and to 150˚C to evaporate all the residual water. Finally, the solution was heated to 300˚C in an argon atmosphere and kept at this temperature for 90 min for complete reaction and crystal formation. After reaction and cooling down to room temperature, the synthesized nanoparticles were washed with cyclohexane/ethanol for several times and dispersed in cyclohexane for use. Other $NaYF_4$: 4% $Tm^{3+}$, $x$% $Yb^{3+}$ ($x$= 30, 45), $NaYF_4$: 8% $Tm^{3+}$, $y$% $Yb^{3+}$ ($y$= 20, 40, 60) samples were synthesized as the same route using varied concentrations.

#### 3. Synthesis of hexagonal phase core-shell structure nanoparticles:

For the synthesis of core-shell structure nanoparticles, typically, $NaYF_4$: 4% $Tm^{3+}$, 20% $Yb^{3+}$ @ $NaYF_4$ nanoparticles. A modified hot-injection method was used for growing shells onto the core nanoparticles. 0.2 mmol $NaYF_4$: 4% $Tm^{3+}$, 20% $Yb^{3+}$ nanoparticles were dispersed in cyclohexane and mixed with OA (3mL) and ODE (8mL) in a 50mL three-neck flask. The mixture was degassed under Ar flow and kept at 100˚C for 30 min to completely remove cyclohexane. Then it is heated up to 150˚C for 15 min to remove water. The mixture solution was then quickly heated to 300˚C and $NaYF_4$ source solution was injected into the core nanoparticles mixture solution using a syringe. The injection rate is 0.05 mL/2 min. After the reaction, the precipitate was washed with cyclohexane/ethanol for several times and dispersed in cyclohexane for use. Other samples, such as $NaYF_4$: 4% $Tm^{3+}$, $x$% $Yb^{3+}$ ($x$= 30, 45) @ $NaYF_4$, $NaYF_4$: 8% $Tm^{3+}$, $y$% $Yb^{3+}$ ($y$= 20, 40, 60) @ $NaYF_4$, were synthesized as the same route.

**4, Monodispersed single UCNPs samples**

1, Place a drop of 50 µL Poly-L-lysine solution (0.1% w/v in $H_2O$) on a cleaned cover-glass and leave it for 30 mins before rinse it using water and dry the surface at room temperature.
2. Prepare 0.01 mg/mL upconversion nanoparticles (UCNPs) dispersion in cyclohexane and place a drop of 20 µL UCNPs dispersion onto the cover-glass, then carefully rinse it using cyclohexane and let it dry naturally.
3. Make a drop of 20 µL Embedding Media on a glass slide, and then place the cover-glass onto the glass slide with Embedding media and squeeze out any air bubbles. Then dry the sample in an oven at 60˚C.

**Characterizations:**

**1. TEM Characterization:**

The morphology of the synthesized nanoparticles were characterized using transmission electron microscopy (TEM) imaging (Philips CM10 TEM) with an operating voltage of 100 kV. The samples were prepared by placing a drop of a dilute suspension of nanoparticles onto the formvar-coated copper grids (300 meshes) and allowing it to dry in a desiccator at room temperature.

**2. Photoluminescence characterization of single UCNPs using laser Scanning Microscopy:**

During the point by point scanning process, when the excitation laser beam moves closer to a single nanoparticle, the system will detect a brighter emission intensity. Therefore, each of single nanoparticle will present a Gaussian spot in the large field PMT-based laser scanning microscope image. The maximum brightness value (photon count) of each Gaussian spot can be used to represent the brightness of that single nanoparticle. For each image, we record all of the single nanoparticles' brightness values (photon counts). For each batch of samples, we average more than 15 single nanoparticles' values to give a mean brightness with standard deviation as error bars.

**Supporting results:**

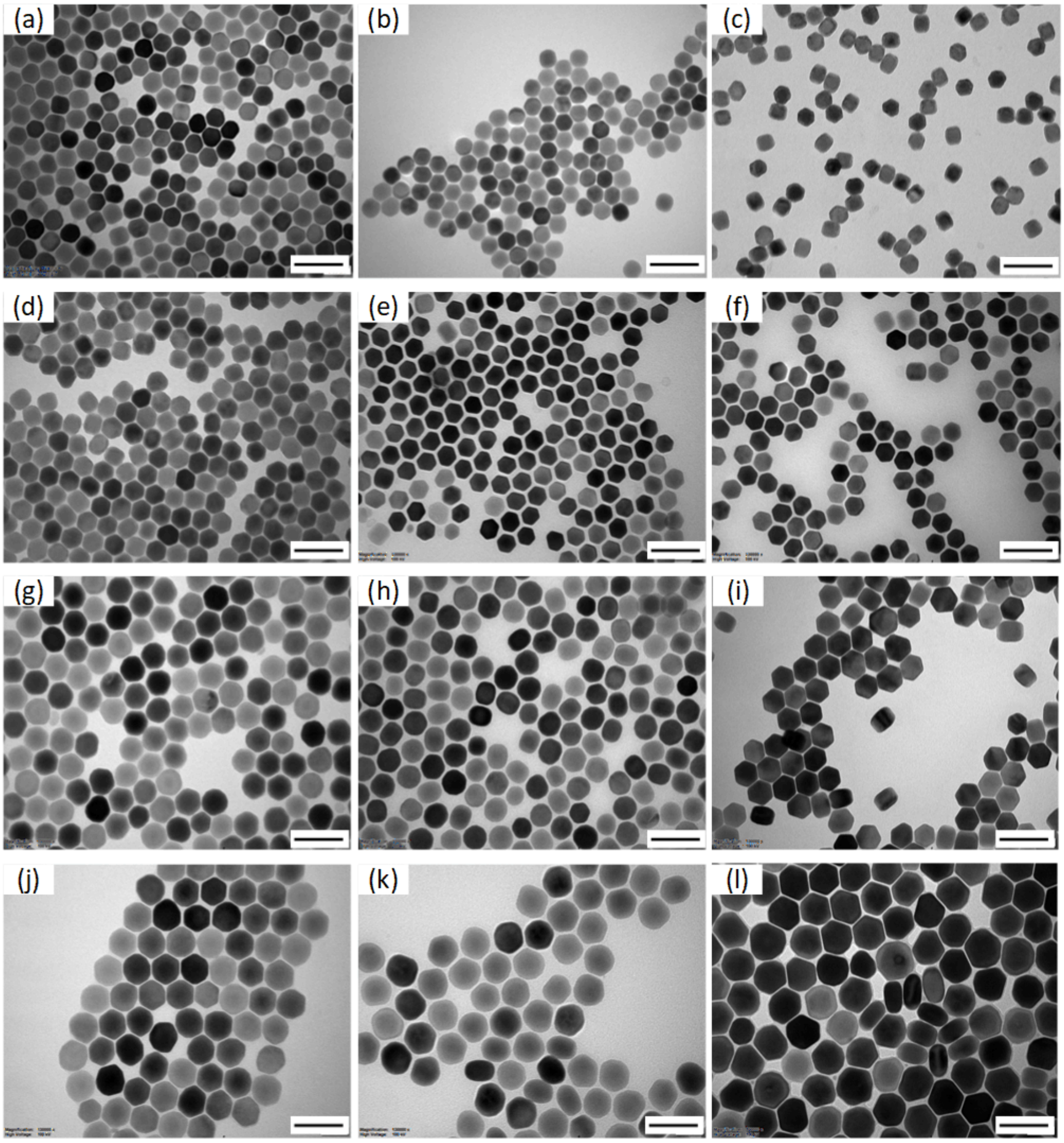


*Figure S1 TEM images of $NaYF_4$:(a) 4% $Tm^{3+}$, 20% $Yb^{3+}$, (b) 4% $Tm^{3+}$, 30% $Yb^{3+}$, (c) 4% $Tm^{3+}$, 45% $Yb^{3+}$, (d) 4% $Tm^{3+}$, 20% $Yb^{3+}$@$NaYF_4$, (e) 4% $Tm^{3+}$, 30% $Yb^{3+}$ $NaYF_4$, (f) 4% $Tm^{3+}$, 45% $Yb^{3+}$@$NaYF_4$, (g) 8% $Tm^{3+}$, 20% $Yb^{3+}$, (h)8% $Tm^{3+}$, 40% $Yb^{3+}$, (i) 8% $Tm^{3+}$, 60% $Yb^{3+}$, (j) 8% $Tm^{3+}$, 20% $Yb^{3+}$@$NaYF_4$ (k) 8% $Tm^{3+}$, 40% $Yb^{3+}$@$NaYF_4$, (l) 8% $Tm^{3+}$, 60% $Yb^{3+}$@$NaYF_4$ UCNPs.*

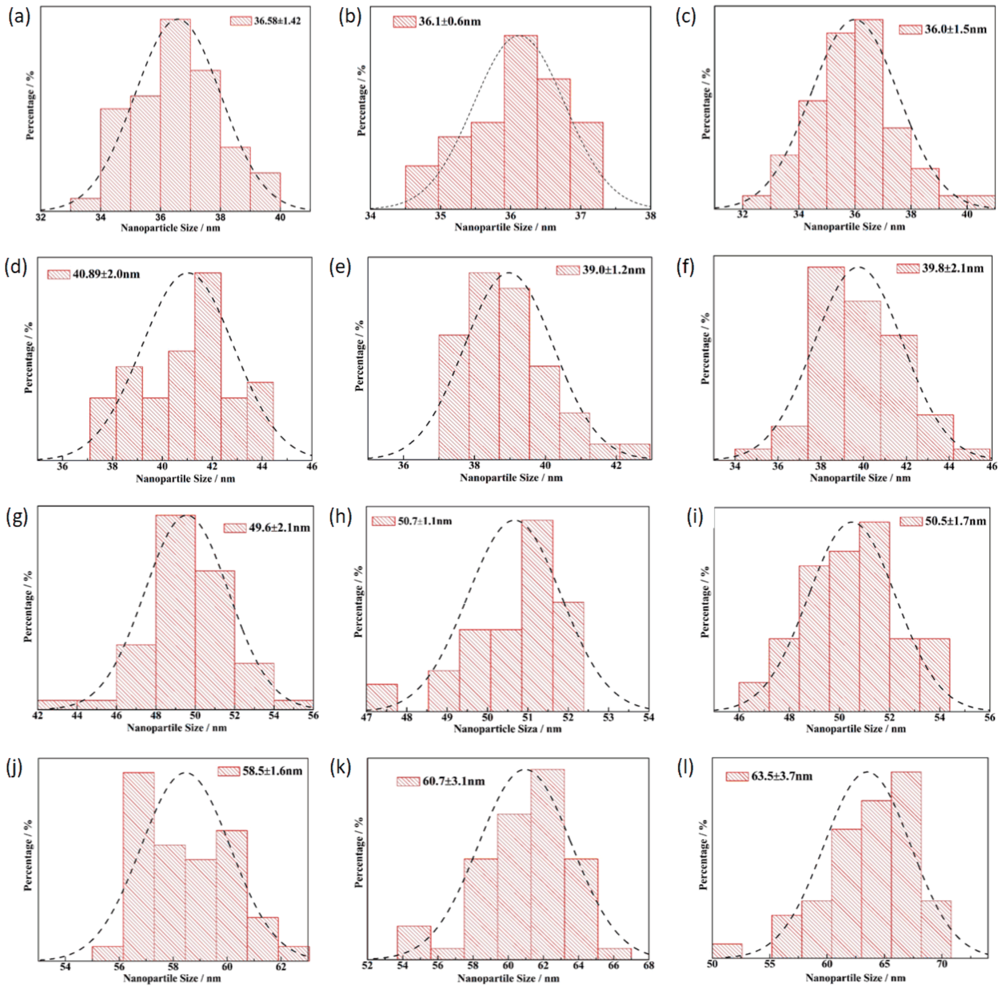


*Figure S2 The nanocrystal size distribution histograms corresponding to TEM images in Figure S1. Histograms of the crystals sizes are drawn from analysis of >150 crystals for each sample. The mean and standard deviation for each monocrystalline diameter are (a)36.6±1.4 nm; (b)36.1±0.6 nm; (c)36. ±1.5 nm; (d)40.9±2.0 nm; (e)39.0±1.2 nm;(f)39.8±2.1 nm; (g)49.6±2.1 nm; (h)50.7±1.1 nm; (i)50.5±1.7; (j)58.5±1.6 nm; (k)60.7±3.1 nm; (l)63.5±3.7 nm.*
